# Cancer Immunotherapy through Tissue Adhering Polymers

**DOI:** 10.1101/2023.03.23.533909

**Authors:** Neil J. Borthwick, Caitlin L. Maikawa, Sven Weller, Thomas L. Andresen, Anders E. Hansen, Anton A.A. Autzen

## Abstract

TLR 7/8 agonists are highly potent immunostimulators, though their clinical translation has been met with mixed success, due to their high toxicity as a result of an unregulated systemic immune activation. There is enormous potential to augment cancer immunotherapies with synthetic TLR 7/8 agonists, though a thorough control of pharmacokinetics and localization is needed for the general use of TLR 7/8 agonists in cancer immunotherapy. Herein, we control localization of TLR 7/8 agonists, by exploiting the extensive tissue retention of poly(acrylic acid-co-styrene). In a murine CT26 model, we find that covalently attaching TLR 7/8 agonists to the copolymer allows for retaining the drug in the tumor microenvironment for at least 15 weeks, after intratumoral injection, and results in a curative monotherapy. The copolymer itself is a new avenue for attaining prolonged tissue rentention for covalently attached drugs.

## Introduction

Cancer immunotherapies have developed as research into the immune system has shed light on mechanisms of tumor cell evasion mechanisms.^1^ Immunotherapies can use antibodies to inhibit immune checkpoints, which in turn enhances an immune response. ^2^ Consequently, immune checkpoint inhibition has proven to be an effective front line treatment against many cancers, with marked effect on cancers with a high mutational burden.^3^ However, the overall response of these therapies varies, and complementary immune therapies are warranted.^3–5^ The PD-L1 pathway directly inhibits cytotoxic CD8 T-cells, as they attack the tumor, and Antibodies against the PD-1 receptor and the PD-L1 ligand have shown increased efficacy pre-clinically if combined with other immune system-stimulating drugs, including Toll-Like Receptor (TLR) agonists, and Stimulator of Interferon Genes (STING) agonists.^6, 7^ The combination of multiple avenues of immune activation shows promise towards enabling a curative response against cancers resistant to checkpoint inhibitors.^8^ The most explored synthetic TLR agonists, are agonists for TLR 7 and 8. These TLR receptors are expressed by most immune cells, and are stimulated by single-stranded RNA. Their activation leads to the production of proinflammatory cytokines, and type I interferons, dependent on the specific cell type.^9^ The TLR 7 agonist Imiquimod, is the only FDA approved TLR agonist for use in cancer immunotherapy. Imiquimod is applied topically to treat melanomas and basal cell carcinomas. More potent TLR 7/8 agonists have been developed, such as Resiquimod, though systemic exposure results in severe side effects. As such, the major challenge of TLR 7/8 agonists is their toxicity. ^10, 11^ The broader use of TLR 7/8 agonists in immunotherapies will require controlled pharmacokinetics, to reduce off-target effects and systemic toxicity. ^12^ Limiting their effects to the injection site can expand the applicability of TLR 7/8 agonists. Moreover, their propensity to stimulate a Th1 response makes them attractive candidates for vaccine adjuvants. The expanded applicability of synthetic TLR 7/8 agonists have been established by several studies. Polymers carrying TLR 7/8 agonists have been shown to be powerful adjuvants in vaccines, while also stimulating class-switching of IgG antibodies, indicative of a Th1 polarized response, towards a cytotoxic cell-mediated immune response.^13, 14^ This approach has also been applied to conjugate vaccines of TLR 7/8 agonists, creating potent personal vaccines in cancer immunotherapy.^15^ Additionally, immunogenic polymeric conjugates or supramolecular assemblies can be injected intratumorally, or peritumorally, and potentiate anti-PD-1, IL-2, and potentially many other complementary immune therapies.^16–19^ Intratumoral and peritumoral injection circumvents the pharmacokinetic challenges, by depositing a high concentration of the TLR 7/8 agonists, directly at the tumor, where it in turn will activate immune cells that are already exposed to tumor antigens. The limited exposure to immune cells systemically also decreases off-target side effects and toxicity.^16, 18, 20^ While these local exposure, and local depot effect are effective, the approach can be improved through a combination of tissue-adhesion, and tumor penetration. A consistent presentation of TLR 7/8 agonists within the tumor could effectively remodel both the tumor environment, and host response, towards an improved clinical outcome.^21^ In this work, we show that amphiphilic polymers can be used to improve the pharmacokinetic properties of TLR 7/8 agonists, with the result being an improved therapy.

Amphiphilic polymers have a long history in drug delivery. Notably, poly(styrene-co-maleic acid) (SMA) was found to have extended circulation upon intravenous injection, with the copolymer being well-tolerated in clinic.^22, 23^ The Enhanced-Permeation and Retention effect (EPR) was first found with this polymer.^24^ The properties of SMA are highly dependent on the ratio of styrene to maleic acid along the polymer backbone, the gradient of this composition along the backbone, as well as the molecular weight.^25^ These polymers are of interest within molecular biology, as they have been found to solubilize bilayer lipids into nanoscale lipid-polymer nanodisc assemblies.^26–29^ We have previously optimized the structurally similar polymer poly(acrylic acid-co-styrene) (AASTY) for the solubilization of membrane proteins into hybrid polymer-lipid bilayer nanodiscs. ^26^ Herein we find that AASTY is retained in tissues for an extended period of time upon subcutaneous injection, despite being a low molecular weight, water soluble polymer, and thus can be useful as a tissue-residing drug depot. We imagine that the polymer associates with lipid bilayers, and at the surfaces of components of the extracellular matrix, slowing lymphatic drainage. Combining this feature with covalently attached TLR 7/8 agonists presents and opportunity for a consistent activation of immune cells at the site of injection, enabling the use of TLR 7/8 agonists in the treatment of solid tumors, while limiting off-target effects, and systemic side effects.

## Experimental

### Reagents

The Cyanine7-DBCO and sulfo-cyanine7.5-maleimide dyes were purchased from Lumiprobe. 4-Amino-2-(ethoxymethyl)-1H-imidazo[4,5-c]quinoline-1-propanamine was purchased from Angene chemicals. All other chemicals were purchased from Sigma-Aldrich or VWR chemicals.

### Synthesis and characterization of AASTY copolymers

The synthesis of the AASTY copolymers was performed as in Smith et al. ^26^ and Timcenko et al..^27^ In short, the copolymers were synthesised using either the RAFT Agent 2-cyano 2-propyl dodecyl trithiocarbonate (1.10 g, 3.17 mmol; giving the copolymer AASTY_12.5_) or 2-(Dodecylthiocarbonothioylthio)-2-methylpropionic acid 3-azido-1-propanol ester (0.052 g, 0.12 mmol; giving the copolymer AASTY_9.9_-N_3_) in Schlenk flasks. The initiator azobisisobutyrunitrile (AIBN) and the monomers acrylic acid (AA) and styrene (STY) were added to the flasks (for AASTY_12.5_: AIBN: 104 mg, 0.634 mmol, AA: 13.0 g, 190 mmol, STY: 26.6 g, 232 mmol; for AASTY_9.9_-N_3_: AIBN: 3.8 mg, 0.023 mmol, AA: 0.324 g, 4.50 mmol, STY: 0.573 g, 5.50 mmol). Oxygen was removed via 4 freeze-pump-thaw cycles. The mixtures were heated to 60 or 70 °C for 9 to 12 hours, reaching 95% monomer conversion for AASTY_12.5_ and 69% for AASTY_9.9_-N_3_. The products were solubilized in diethyl ether, precipitated into heptane, and dried in vacuo to yield yellow crisp solids. For AASTY_12.5_, the polymer was solubilized in (water:ethanol)(1:3) with 30% H_2_O_2_ and incubated at 70 °C overnight to remove the dodecyl trithiocarbonate end group (ttc), yeilding a colorless solution. That polymer was precipitated into deionized water and collected by centrifugation. Final polymer products were converted to partial sodium salts by solubilization in deionized water with the addition of NaOH (1 M) until the pH was at 7-7.5, and the opaque mixtures were filtered and lyophilized. The characterization of the polymers was performed as in Smith et al.,^26^ ^1^H-Nuclear Magnetic Resonance (NMR) was used to measure the monomer conversion using a Bruker Avance III 400 MHz system. The number average molecular weights (M*_n_*), weight average molecular weights (M*_w_*) and dispersities (D = M*_n_*/M*_w_*) were measured using a Dionex Ultimate 3000 system. Detection was performed by a Dawn Heleos II Multi Angle Light Scattering detector, and an Optilab rEX refractive index detector. Gel Permeation Chromatography (GPC) was done in a Superose 6 Increase column (10/300, Cytiva). Data were analyzed with Astra 7.0, using a dn/dc of 0.170 mL/g.

### Polymer conjugation with TLR agonists and imaging dyes

The AASTY copolymers were added to 5, 10 or 15% molar equivalent to the AA content of the TLR 7/8 agonist 4-Amino-2-(ethoxymethyl)-1H-imidazo[4,5-c]quinoline-1-propanamine (refered to as TLR7/8a) in Dimethylformamide (DMF). 2 molar equivalents of N-Hydroxysuccinimide (NHS) with 1 molar equivalent of N,N’-Diisopropylcarbodiimide were added to the samples, with respect to the TLR7/8a. The reactions were left at room temperature overnight, and the products were purified the next day by a NAP^TM^ DNA purification column, and lyophilized to yield white solids. They were finally characterized by ^1^H NMR to quantify the extent of conjugation.

A fraction of the TLR 7/8 conjugated AASTY_9.9_-N_3_ or the unmodified AASTY_9.9_-N_3_ were solubilized in methanol with 1 molar equivalent of a Cyanine 7 Near-Infrared dye derivative featuring a dibenzocyclooctyne (DBCO) moiety for copper-free click-chemistry, with respect to end groups. The reactions were left at room temperature overnight, purified the next day by a NAP^TM^ DNA purification column and lyophilized, to yield dark green solids.

### In vitro TLR response to AASTY-TLR7/8a conjugates

The *in vitro* activity of the TLR constructs was verified using commercially available RAW Blue Cells (InvivoGen Cat raw-sp, LOT R01-4301) according to the manufacturer’s protocol with minor modifications. Briefly, cells were cultured up to 80% confluency in T-25 culture flasks in Dulbecco’s modified Eagle’s medium (DMEM), containing 20% fetal bovine serum. 2 mM L-glutamine, 4,5 g/l glucose, 100 *µ*g/ml Normocin and 1% Pen-Strep. The selection antibiotic Zeocin was added every third passage to maintain the selection pressure. Three days before the assay, cells were seeded at a concentration of 1.5×10^4^ per cm^2^ and collected with a cell scraper at a final concentration of 550.000 cells/ml on the day of the assay. Off the stock solution, 180 µl were added to flat bottom 96-well plates containing 20 µl of the construct and control TLR 7/8 agonist respectively. After 24 h incubation time, 20 µl of the induced RAW-Blue cells supernatant was mixed with 180 µl QUANTI-Blue solution. The absorbance of the plate at 620 nm was measured after incubating for 5 h 30 min at 37°C with a TECAN spark plate reader.

### Animal experiments

The experimental procedures were all internally reviewed to be in accordance with the license approved by the Danish National Animal Experiment Inspectorate.

#### Biodistribution studies

BALB/cJRj mice were injected with 100 *µ*L of (AASTY_9.9_-Cy7:AASTY_12.5_) (1:10) in Dulbecco’s phosphate-buffered saline (DPBS) at 1 mg/mL, for an absolute dose of 1.5 nmol Cy7, either subcutanously on the left flank or in the tail vein. Controls received an equivalent dose of the free dye in a sulfonated water-soluble form (sulfo-Cyanine7.5 maleimide), defined by equal absorbance at 710 nm. At regular intervals, they were then anaesthetized by isoflurane inhalation and imaged in a MILabs U-CT Optical Imaging instrument in fluorescence mode (*λ_ex_* = 710 nm; *λ_em_* = 785 nm). The processed image files were then analyzed with ImageJ 1.47t. Signal quantification was performed by measuring the integrated density on Regions Of Interest (ROI) automatically delimited by Blob detection on grayscale images. The exponential decay of the Cy7 signal in these ROIs allowed to calculate tissue-specific half-lives for the Cy7 conjugates. After the last time-point, the mice were euthanized, dissected to extract the liver, spleen, kidney and inguinal lymph node on the injection side. Acquisitions were made on the organs and the dissected corpse to screen for residual signal, and here the ROIs were delimited manually due to the poor signal-to-noise ratio. During the whole course of the study, the mice were also monitored for signs of misthriving, visible tissue damage or inflammation at injection site, none of which was observed.

#### Cancer studies

BALB/cJRj mice were injected with 100 *µ*L of Murine colon carcinoma CT26 cancer cells (3×10^5^) in serum-free Roswell Park Memorial Institute (RPMI) medium subcutanously on the right flank. Mice bearing tumors with volumes between 50 and 200 mm^3^ were assigned to one of four treatments (See Table 1) using a randomized block design based on initial tumor volume. Mice were rearranged into new cages where each cage received one of the four treatments. Treatments were administered intratumorally (50 *µ*L). A total of 3 injections, separated by 7 days each, were made for each treatment.

**Table 1:** Summary of tested treatments

| Treatment name | Compound | Concentration (mg/mL) | buffer | TLR7/8a dose (mg/kg) | Cy7 dose (nmol) |
| --- | --- | --- | --- | --- | --- |
| free TLR7/8a | TLR7/8a | 2.1 | 3.5% DMSO-DPBS | 5 | 0 |
| AASTY-Cy7 | (AASTY<sub>9,9</sub>:AASTY<sub>9,9</sub>-Cy7) (1:90) | 16.0 | DPBS | 0 | 1.5 |
| AASTY-TLR7/8a-Cy7 0.5X | (AASTY<sub>9,9</sub>-TLR7/8a-Cy7:AASTY<sub>9,9</sub>-TLR7/8a) (1:45) | 9.2 | DPBS | 2.5 | 1.5 |
| AASTY-TLR7/8a-Cy7 1X | (AASTY<sub>9,9</sub>-TLR7/8a-Cy7:AASTY<sub>9,9</sub>-TLR7/8a) (1:90) | 18.4 | DPBS | 5 | 1.5 |

83 days after the first tumor inoculation, the surviving mice were re-challenged with a second inoculation of CT26 cancers cells, using the same dose, on their left flank. A new control group of the mice received their first inoculation of cells at a same cell dose as previously. Mice were monitored twice a week. At these time-points, they were weighed and tumor volumes were measured with electronic calipers. Fluorescence imaging was performed 7 min, 21 h, 48 h and once a week after the first injection for all formulations containing Cy7 moieties. Once euthanized, the mice were stored at -20 °C and autopsied within one week in the same way as for the previous biodistribution studies but also including both inguinal lymph nodes and the resected tumors. Considering that these mice were euthanized at different time-points during the study, relative signal quantification was preferred here and was calculated by dividing the integrated signal of each organ by the total integrated signal per mouse.

The biodistribution of the compound was further assessed by injecting a single dose of AASTY-TLR7/8a-Cy7 1× (50 *µ*L, intratumor) and scanning the mice by Fluorescence Tomography (MILabs U-CT in FLT mode, *λ_ex_* = 730 nm; *λ_em_* = 785 nm). Due to a maintenance of the equipment, the scans were performed 4 to 5 days after euthanasia of the mice, which were stored at -20 °C. The reconstructed images were analyzed and 3D-rendered using Imalytics Preclinical 2.1.

During the study, tumor volumes were calculated as length x width^2^ / 2 and the mice were euthanized when tumor volumes reached 2000 mm^3^. Humane endpoints included general signs of misthriving mice, significant weight loss (*>*15% of initial weight or *>*10% overnight weight loss), or presence of wounds larger than 8 mm on the tumors.

### Statistics

Statistical analyses were performed on SAS 9.4. To meet the assumption of homoscedasticity, analysis of mean tumor growth was only performed for the first 10 days of the treatment and the tumor volumes were transformed using natural logarithm. The values were then analyzed using a restricted maximum likelihood (REML) mixed model, with mouse included as random effect subject. To assess whether treatment altered tumor growth over time, the interaction between treatment and time was measured, and the regression slopes of each treatment were then compared post-hoc pairwise using a Bonferroni correction to adjust for multiple comparisons (*α* = 0.005). Analysis of survival was performed using a maximum likelihood parametric regression with censored data assuming a Weibull distribution. The survival times of the different treatment groups were compared through their predicted least-squared means and Tukey-Kramer post-hoc tests were used to adjust for multiple comparisons. A Kaplan-Meier plot has been included for data visualization, but no additional statistical testing was performed. For the rest of the plots, all data are presented as the means ± Standard Error of the Mean (SEM) unless stated differently. Statistical significance was defined as *α <* 0.05 where not otherwise specified.

### Note

No unexpected or unusually high safety hazards were encountered during the course of this work.

## Results and discussion

### AASTY can be easily functionalized while retaining its properties

Two AASTY polymers were tested to serve as the backbone for functional conjugations: AASTY_12.5_ (12.5 kDa), developed and tested by Smith et al. ^26^ and Timcenko et al., ^27^ and AASTY_9.9_-N_3_ (9.9 kDa, with an Azide moiety), developed specifically for this experiment. They were selected for their small size lying below the glomerular filtration threshold,^30^ and their characteristics are summarized in Table 2.

**Table 2:** Summary of polymer characteristics

| Copolymer | initial AA content | % conversion | M<sub>n</sub> (kDa) | D |
| --- | --- | --- | --- | --- |
| AASTY<sub>12.5</sub> | 0.45 | 0.95 | 12.5 | 1.1 |
| AASTY<sub>9.9</sub>-N<sub>3</sub> | 0.45 | 0.69 | 9.9 | 1.4 |

Using copper-free click chemistry, the azide moiety on the R terminus of AASTY_9.9_-N_3_ was used to attach a Cyanine 7 Near-Infrared dye for tracking the copolymer in biodistribution studies. In a AASTY_9.9_-Cy7:AASTY_12.5_ (1:10) formulation, the dye did not affect the nanodisc-forming properties of AASTY, measured through solubilization of lipid small unilamellar vesicles (SUV) (data not shown).

The pendant groups of the copolymer, the acrylic acid groups, were used to conjugate a TLR 7/8 agonist to trigger immune responses *in vivo*. We aimed to determine the amount of agonist that could be attached to the copolymer without rendering it insoluble in aqueous media. The extent of conjugation was measured by ^1^H NMR (Fig.S1). We observed that the copolymer conjugate remained soluble in water with up to at least 11% of carboxylates being functionalized with the TLR 7/8 agonist, despite the hydrophobicity of the agonist and the high content of styrene. The conjugation site on the TLR 7/8 agonist was an amine at a site shown not to interfere with TLR 7/8 receptor binding and activation.^31, 32^ Finally, to assess the TLR stimulation potential, we used the RAW Blue reporter assay and showed that the cellular response from these conjugates was increasing with the TLR 7/8 conjugation (Fig.2.C). The highest conjugation ratio (11.2%) was therefore selected for the cancer studies.

**Figure 1:**
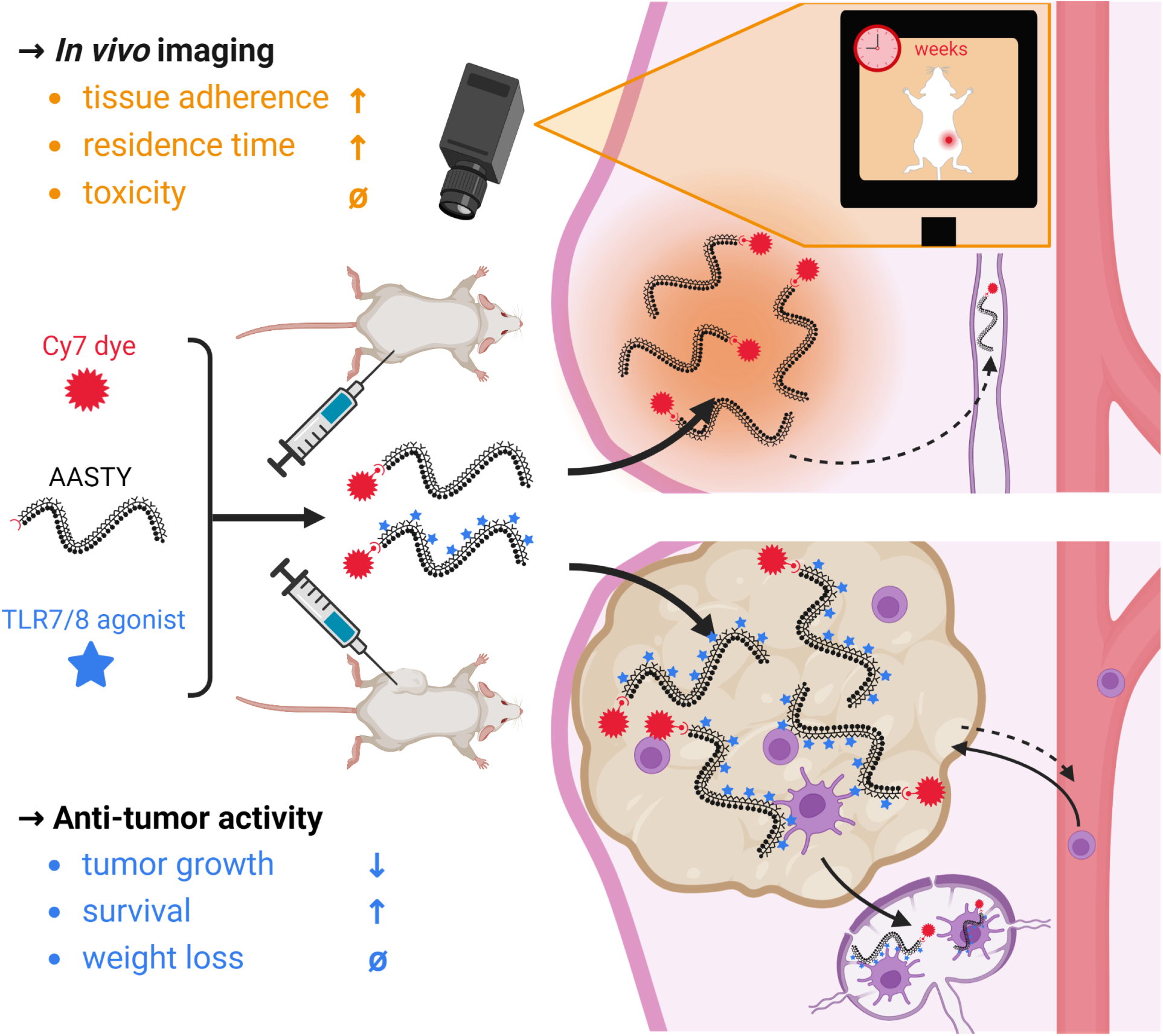
Administration of AASTY-TLR7/8a-Cy7 conjugates in mice for *in vivo* imaging and cancer treatment. AASTY strongly adheres to tissues upon injection. Conjugated to a Cy7 dye, it can be used as a stain for imaging applications for durations ranging from weeks to months. Conjugated to Toll-Like Receptor 7/8 agonists, it can trigger immune responses in the local tumor environment and the lymphatic system to induce tumor elimination. Its ability to adhere to tissues protects the mice against the life-threatening side effects induced by systemic circulation of TLR 7/8 agonists, and overall the treatment results in increased survival.

**Figure 2:**
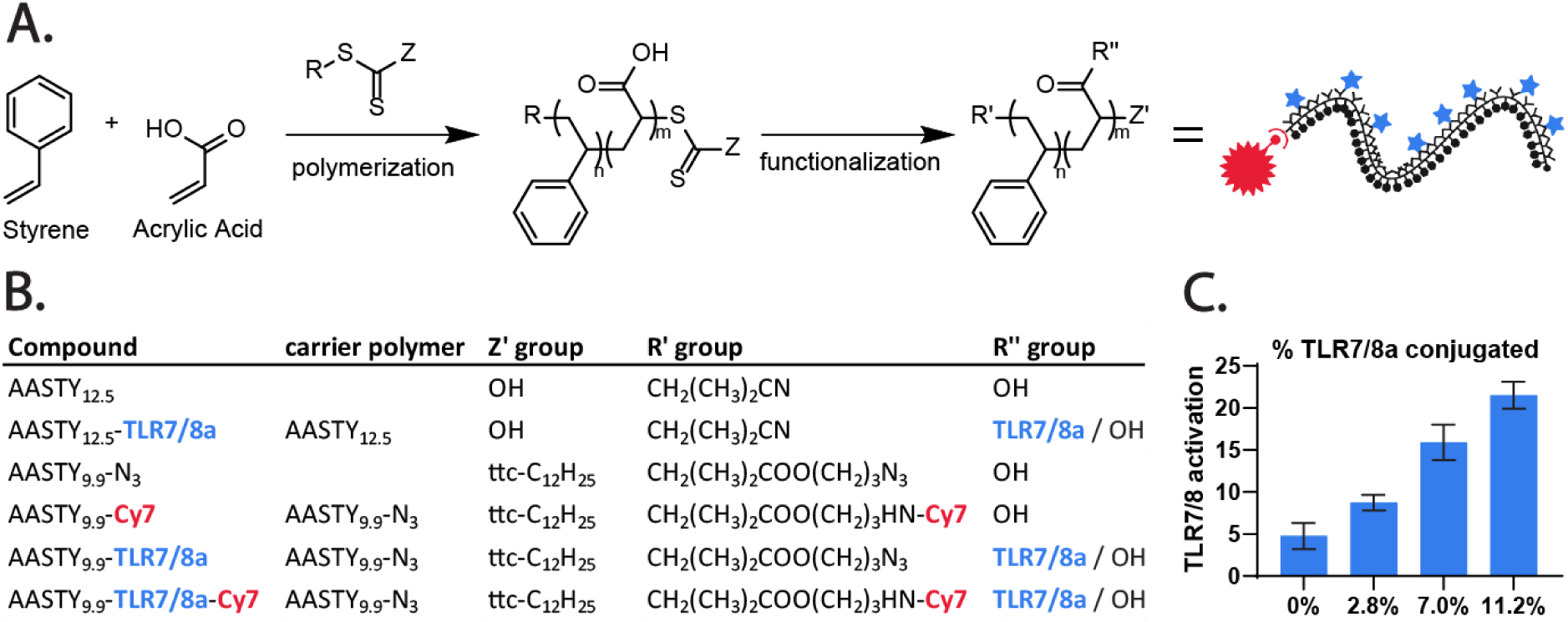
Summary of conjugates characteristics. (A,B) Styrene and acrylic acid are copolymerized using the RAFT technique and using a chain-transfer agent which incorporates functional azide endgroups on the final product. Using azide-alkyne click chemistry, a Cy7 Near-Infrared dye moiety can be conjugated at the R terminus of the polymer, and using carbodiimide coupling, a potent TLR 7/8 agonist can be conjugated on a fraction of the acrylic acid monomers. Optionally, the trithiocarbonate on the Z terminus can also be removed. (C) TLR *in vitro* activation potential of the polymers with increasing extent of TLR7/8a conjugation. Activation was quantified using a RAW-Blue murine macrophage reporter cell assay by measuring absorbance at 620 nm.

To study the biodistribution of AASTY inside tumors during treatment, we successfully combined both approaches described above by conjugating the TLR 7/8a on the AA of AASTY_9.9_-N_3_, up to 10% for that batch, followed by an addition of the Cy7 dye on azide end-group. All the constructs that were developed and used in this study are summarized in Fig.2.A and B.

### AASTY constructs reside in tissues for an exceptional duration *in* ***vivo***

To the best of our knowledge, AASTY copolymers have not been used *in vivo*, their main applications being oriented towards structural biology. Here, for the first time, we administered AASTY constructs (AASTY-Cy7) in mice via the subcutaneous or intravenous routes and monitored their biodistribution over the course of 2 weeks.

This study extended the pharmacokinetics (PK) of the AASTY-Cy7 conjugate following subcutaneous (SC) injection. Results indicated that the majority of the fluorescence signal remained in the vicinity of the initial injection site and did not significantly diffuse away after the formulation was absorbed by the SC tissue (Fig.3.A and S2). While the integrated signal density decreased over time, 13% of the initial signal was still visible at the injection site after 2 weeks, compared to *≺* 1% for the control. This suggests that the AASTY-Cy7 conjugate strongly and rapidly adheres to local tissues for extended periods of time. Image analysis estimated the compound’s half-life in the SC tissue to be 58 hours, which is remarkable for water-soluble material (Fig.3.C and D).^30^ As an additional control, an equivalent dose of the non-water soluble DBCO-Cy7 dye in the form used for conjugation was solubilized in 5% Dimethyl sulfoxide (DMSO)-DPBS and injected SC in a similar manner. The control experiment showed that despite not being cleared through renal excretion, the free dye is rapidly metabolized at its injection site with no visible signal remaining after 1 week, which confirms that the conjugates exhibit different PK profiles than the free forms of the dye (Fig S3). Water soluble sulphonated Cy7.5 (”free Cy7”) was rapidly cleared by renal elimination (*≺* 2% remaining after 24h) (Fig.3.A and S2). Considering that no significant signal was detected outside the injection region *in vivo*, the mice were autopsied at the end of the 2 weeks to extract organs and analyze the signal distribution with greater sensitivity. The isolated organ images revealed that signal was emanating from a spot on the inguinal lymph node on the injection side, *≈* 11% of the total signal per mouse, and residual signal was also visible in the liver, *≈* 76% (Fig.3.F and S4). This suggests that the compound is slowly cleared via hepatic and lymphatic routes. Based on observations of the living mice during the whole study or of the organs during autopsy, no visible signs of toxicity or inflammation were observed, e.g. swelling, redness, necrosis, misthriving mice, abnormal organ size or aspect, etc., suggesting that AASTY-Cy7 itself doesn’t exhibit high toxicity or immunogenicity.

**Figure 3:**
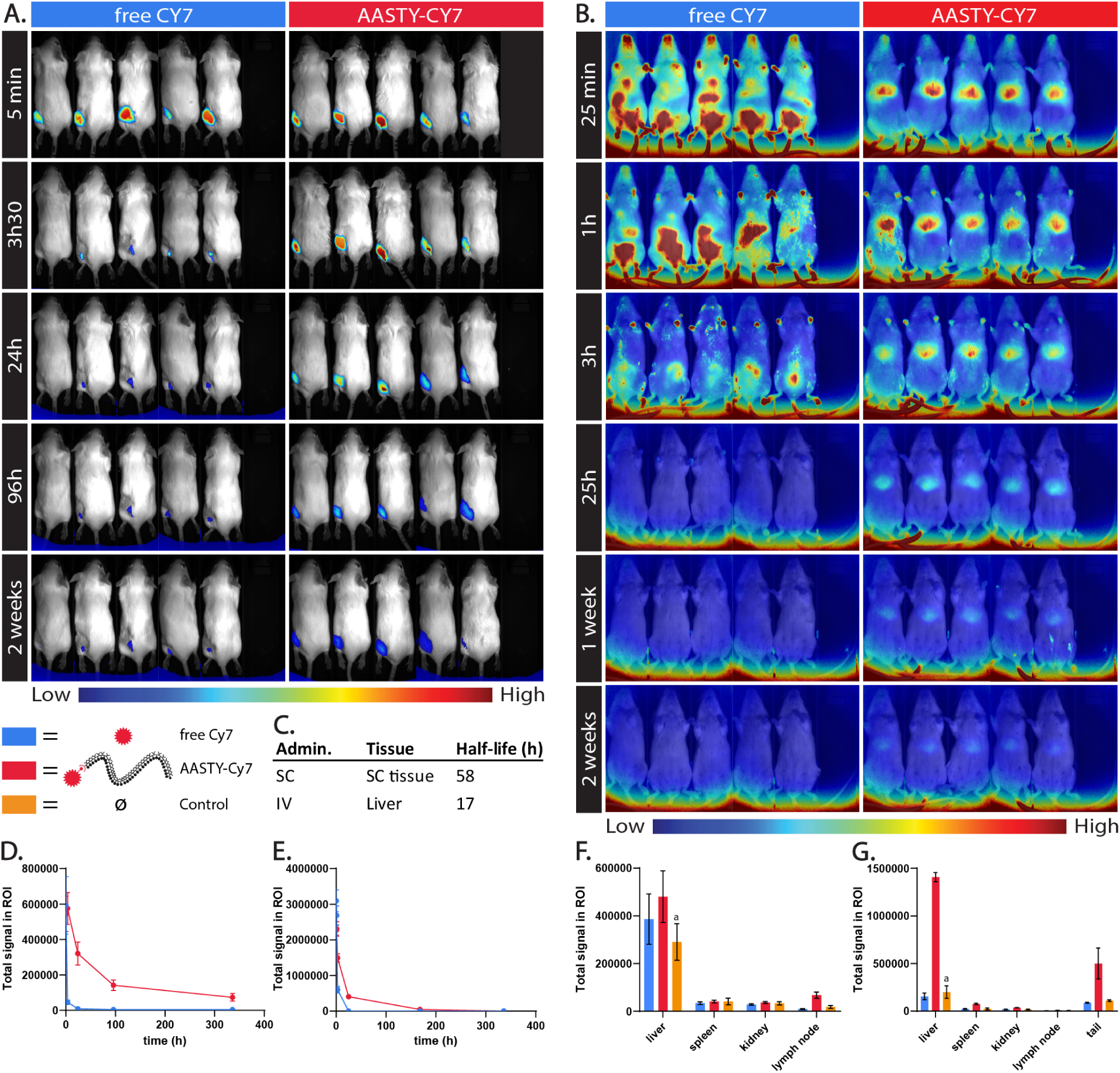
Biodistribution of AASTY-Cy7 conjugates after SC (A,D,F) or IV (B,E,G) injection. (A,B) *In vivo* fluorescence imaging acquisitions of mice after injection of AASTY_9.9_-Cy7 or a water-soluble free Cy7 as a control. (C,D,E) Cy7 signal quantification, based on the unitless integrated density from ImageJ 1.47t, and half-life estimation of the images in (A) and (B) (half-lives were estimated using a linear regression of the natural log-transformed signal intensity). (F,G) Cy7 signal quantification of images from different organs during autopsy 2 weeks after injection. The data are represented as means *±* SEM (n = 5), except for the controls in (F) and (G) (noted with an *^a^*). For these, the error represents the signal SEM of a single mouse oriented in 2 extreme positions to account for the variability brought by the strong background noise at the bottom of the images (visible in (B)).

The next study aimed to investigate the pharmacokinetics of the AASTY-Cy7 conjugate following intravenous (IV) injection in the tail vein. Results indicated that the compound rapidly reached systemic circulation, as evidenced by signal visible in the paws, snout, and eyes, for the first few hours after injection (Fig.3.B). Similar to the SC injection study, a significant portion of the signal remained close to the injection site, in this case, the tail, and did not diffuse for the entire duration of the study. An important signal was visible in the liver from the first acquisition time until the end of the 2 weeks. Image analysis of the liver region revealed a shorter half-life for the compound in the liver, of about 17 hours (Fig.3.C and E.). The actual value is certainly slightly higher considering that the signal of the 2 weeks time-point is below the quantification threshold of the Blob analysis, due to the high noise, but that it was nevertheless still visible on the scans (Fig.3.B). The organ extraction and analysis after autopsy, in which the tail of the mouse was included, confirmed accumulation of the compound in the liver, *≈* 71% of total signal per mouse, including the tail. It also showed a small accumulation in the spleen, *≈* 4%, which was not observed for SC administration (Fig.3.G and S4). No significant signal was reported in the inguinal lymph node or the kidney. Here again, no outward visible signs of toxicity or inflammation were observed.

Together, these two studies suggest that AASTY has the ability to adhere to the local tissues in which it is injected and remains for remarkably long periods of time. Yet the data also suggests that AASTY is slowly cleared over time through the hepatic and lymphatic route. The polymer styrene-co-maleic acid (SMA), a polymer with similar structure to AASTY that was tested *in vivo*, is known to adhere to serum proteins like albumin.^33^

Considering that AASTY can be expected to behave similarly, it would explain how it entered systemic circulation but was not filtered by the kidneys, despite its low molecular weight. Water-soluble polymers are otherwise rapidly eliminated, with slight variations depending on the route of administration.^30^ The control experiment also points out that the dye is likely not cleaved from AASTY during its *in vivo* residency. Adding on top of that, the good tolerance of the mice to the compound and the apparent absence of immunogenicity, this makes AASTY conjugates ideal candidates for retaining a dye at an injection site, or attaining local drug exposure.

### Local exposure to AASTY-TLR7/8a agonist constructs results in an anti-tumor immune response

The AASTY-Cy7-TLR7/8a conjugate was evaluated in a mouse cancer model to assess its potential as a targeted treatment. The study was based on the observations made during the biodistribution studies, which suggested that the polymer’s properties could enable localized and extended exposure of the TLR 7/8a to tumor tissues. This, in turn, was hypothesized to elicit a robust innate and adaptive anti-tumor immune response while minimizing the side effects associated with systemic TLR 7/8a exposure. Additionally, the Cy7 moiety of the conjugate enabled monitoring of the compound’s biodistribution in the tumor and other parts of the body.

The study evaluated the effect of the AASTY-TLR7/8a-Cy7 conjugate on tumor kinetics using mouse cancer models. Tumor growth was monitored and compared across different treatment groups, including control, AASTY-Cy7 alone, TLR7/8a alone, and AASTYTLR7/8a-Cy7 at different doses (Fig.4.C to G). The results showed that most tumors grew rapidly in all groups except for the group treated with the AASTY-TLR7/8a-Cy7 1X dose (Fig.4.G). The influence of two parameters was assessed: the dose of the TLR7/8a, with a non-lethal full dose in the range expected to trigger an immune response and a half-dose below that range, and the dose of AASTY, kept at a constant ratio to the TLR7/8a. A priori, a maximum likelihood parametric regression with censored data assuming a Weibull distribution was chosen to determine if treatment had an effect on survival. Predicted least-squared mean survival time was compared between treatment groups and Tukey-Kramer post-hoc tests were used to adjust for multiple comparisons (Fig.4.K). There was a difference in model predicted-mean survival time between treatments (Wald Chi-Square = 14.6; DF = 4; p = 0.0056). AASTY-TLR7/8a-Cy7 1X had a longer predicted mean survival (118.9 days) compared to the no treatment control (mean = 21.8 days; *|z|* = 3.25; p = 0.010) and AASTY-Cy7 alone (mean = 22.6 days; *|z|* = 3.13; p = 0.015). The absence of response to the AASTY-Cy7 and AASTY-TLR7/8a-Cy7 0.5X treatments suggest that AASTY-Cy7 alone does not exhibit anti-tumor activity. While TLR7/8a alone injected at a full dose intratumorally resulted in two out of six mice completely recovering, no difference in model mean survival time was observed compared to the control (mean = 52.6 days; *|z|* = 2.19; p = 0.18). This is likely because the other four mice did not respond to the treatment. This suggests that the high potency of these compounds may be limited by their rapid diffusion outside the tumor microenvironment after injection. Based on the data from our biodistribution studies, conjugation of TLR7/8a to AASTY-Cy7 will result in sustained local residence times at the tumor site. The AASTY-TLR7/8a-Cy7 0.5X dose did not show improvement in neither predicted mean survival time nor tumor growth rate compared to the control and free TLR7/8a groups. In contrast, the AASTY-TLR7/8a-Cy7 1X dose was the only group to demonstrate an overall negative tumor growth rate in the first 10 days (Fig.4.I, J and S5). For that treatment group, only two mice out of six started to grow tumors after 20 days while all the others completely recovered (Fig.4.G and H). The absence of acute weight loss (*>* 10% overnight or *>* 15% initial weight) or other signs of misthriving mice suggests good tolerance to the treatments (Fig.S6).

**Figure 4:**
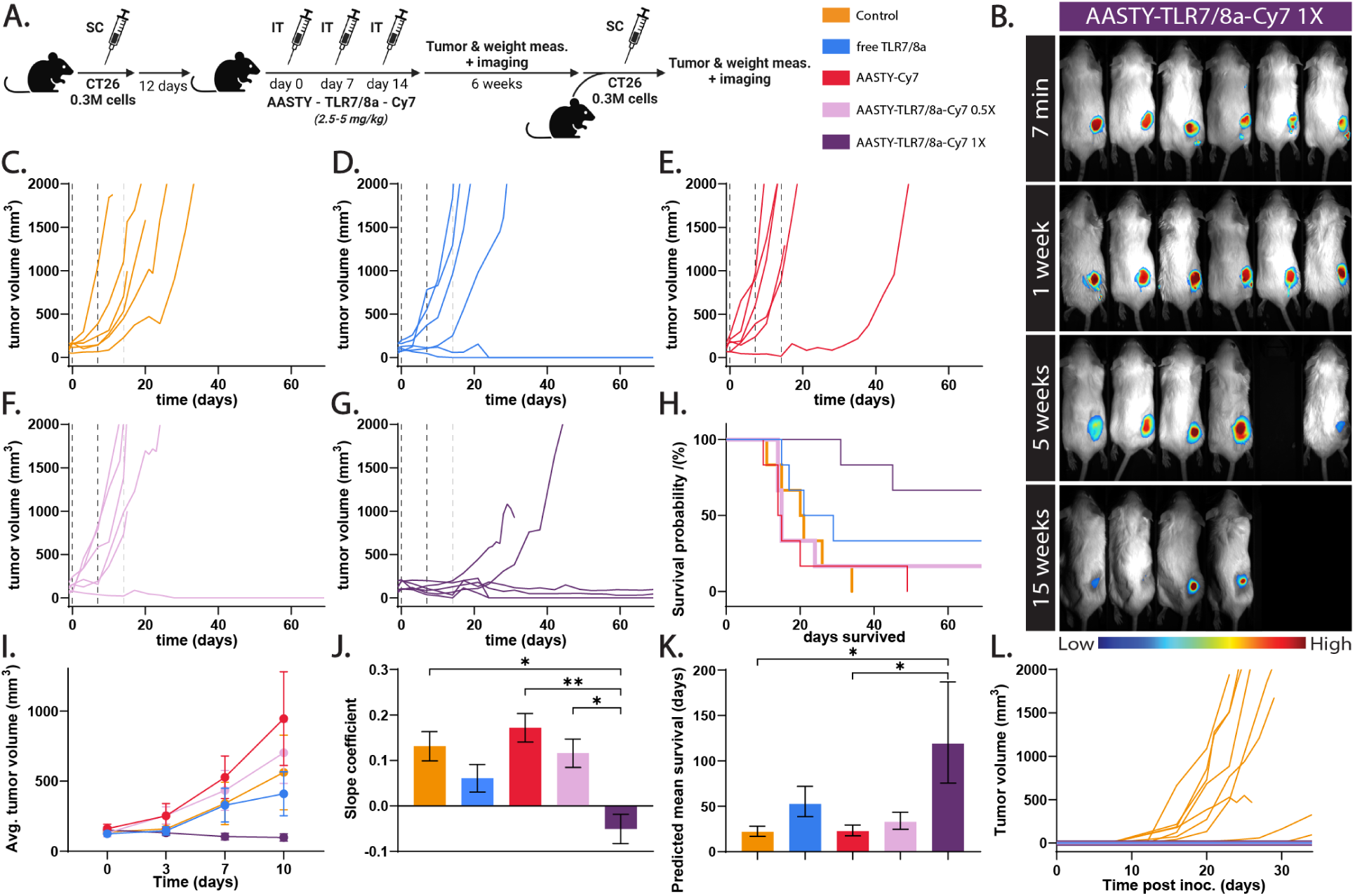
Treatment of Murine Colon Carcinoma (CT26). (A) Study timeline. BALB/c mice were inoculated with 0.3 M CT26 cells SC on the right flank. Mice were treated 12 days later by intratumoral administration 3 times over 2 weeks and tumor size was monitored. Mice were euthanized when tumors reached 1500 mm^3^. After 6 more weeks, the surviving mice were rechallenged with a second CT26 inoculation on the left flank. (B) Fluorescence imaging of the 1X dose mice for the 15 weeks following the first dose injection. (C-G) Individual tumor growth curves for the mice that received: no injection as a control (C), free TLR7/8a treatment (D), AASTY-Cy7 treatment (E), AASTY-TLR7/8a-Cy7 0.5X treatment (half dose) (F), and AASTY-TLR7/8a-Cy7 1X treatment (full dose) (G) (n = 6). The dotted lines represent the injection time-points. (H) Kaplan-Meier survival curves of each group after treatment administration (n = 6). (I) Average tumor growth for each group for the first 10 days following the first dose injection (n = 6). (J) Comparison of the slope coefficients of the log-transformed average tumor growth. (K) Predicted mean survival for each treatment group. (L) Individual tumor growth curves for the surviving mice that were re-challenged with CT26. A new group of mice was included during inoculation as control. (I,J) Data depict mean *±* SEM. (J) The values were analyzed using a restricted maximum likelihood mixed model and the slope coefficients were compared with a Bonferroni-corrected *α* of 0.005 (*p *≺* 0.005, **p *≺* 0.0001). (K) Survival means are shown as inverse link transformed least square means *±* SE predicted by a maximum likelihood parametric regression. Tukey-Kramer post-hoc tests were used to adjust for multiple comparisons (*p *≺* 0.05).

The anti-tumor response observed in the AASTY-TLR7/8a-Cy7 1X dose treatment group was evaluated further to investigate the involvement of an innate and adaptive immune response. The flatness of the growth curves and the immediate aspect of the anti-tumor response observed in 4.G suggest that an innate immune response was involved. To assess the presence of an adaptive response, the surviving mice were re-challenged with tumor cells 83 days after the first inoculation and no tumor growth was observed, supporting the hypothesis that adaptive immune response was involved (Fig.4.L).

In terms of biodistribution, the intratumoral administration data showed an extremely long residence time for the AASTY conjugates, with a half-life in the tumor estimated at 41 days, and 46% of the initial signal was still visible when the study was terminated 91 days after the last dose injection (acquisition week 15) (Fig.4.B and S7). The mice that survived the study didn’t have resected tumors to include in the autopsy so the image analysis of the different organs was separated between ”uncured mice”, including tumors, and ”cured mice”, without. For the uncured mice, *≈* 61% of the total signal was located in the tumor and the rest of the compound mainly distributed in either the liver, *≈* 34%, or the inguinal lymph node neighbouring the tumor,*≈* 3%, (Fig.S8). For the cured mice, the surface signal measured *in vivo* remained in the area neighboring the tumor scar, suggesting that the compound was redistributed in the surrounding tissues as the tumor cells were eliminated. The autopsy of these mice then revealed that *≈* 80% of the remaining compound distributed in the liver, and *≈* 11% in the inguinal lymph node neighbouring the tumor (Fig.S8). Interestingly, for the uncured mice, the relative amount of the the compound in the liver and lymph node increased with the amount of TLR7/8a conjugated to the polymer, which corroborates the idea that phagocytes could play a role in clearance of the TLR7/8a conjugate. However, these mice were euthanized at different stages of their disease, either due to the tumor growing too large or the tumor collapsing, and quantification was done on different group sizes for each treatment since some of the treated mice were cured. Interpretation of the latter set of results should therefore be done carefully.

To gain a better understanding of the biodistribution dynamics after intratumoral administration, fluorescence tomography scans were performed on a small group of extra mice 5 to 6 days after the administration of a single dose of AASTY-TLR7/8a 1X. The results confirmed the previous findings, showing that the compound diffused throughout the entire bean-shaped tumor, with a signal maximum at the injection point, and mostly accumulated in the liver (Fig.5). The new information provided by this study is that the compound apparently distributed throughout the entire lymphatic system, as signal can be detected in a globally symmetric network of canals connected by higher intensity nodes that more or less align with known locations of lymph nodes (notably, among others, the mandibular, cervical, axilliary, brachial, lumbar and codal nodes).

**Figure 5:**
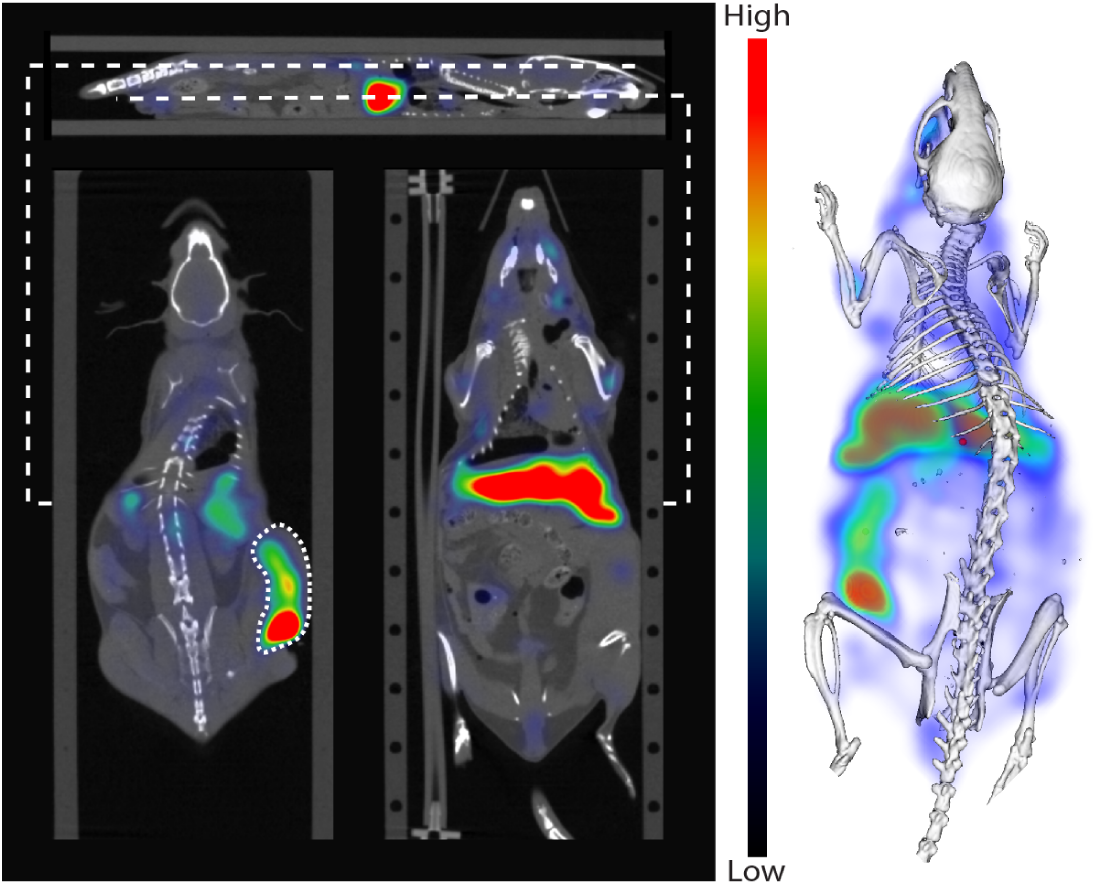
E*x vivo* fluorescence tomography imaging of a CT26 bearing mouse 4 days after injection of a single AASTY-TLR7/8a 1X dose. The Computed Tomography (CT) data is overlayed with the fluorescence signal. Left: one sagittal and two frontal slices of the raw data. The slices are delimited by large white dotted lines, and the tumor on the upper frontal slice is delimited by a small white dotted line. Right: 3D rendering of the data set.

Based on our findings, the following mechanism of action is proposed: upon injection, AASTY rapidly adheres to local tissues, likely by utilizing its amphiphilicity to incorporate itself into lipid membranes. If it is injected intravenously or in a highly vascularized tissue, a fraction of it will flow into the systemic circulation and quickly accumulate primarily in the liver and, to a lesser extent, in the spleen. There is no evidence to suggest that the polymer itself triggers any form of immune reaction. AASTY remains in the injection area for a period ranging from weeks to months but a fraction of it is slowly cleared through the lymphatic system and liver over time. Whether accumulation of the compound in the lymphatic system is mainly driven by systemic circulation or immune cells is still unclear at this point. The pharmacokinetics of AASTY itself cannot be directly determined, as the Cy7 dye used for the biodistribution studies could also potentially be inactivated metabolically. In any case, the extended residence time of AASTY allows for the conjugation of molecules to the polymer, resulting in targeted exposure of specific tissues to the compound. In this way, conjugated TLR 7/8 agonists can be restrained within tumors and locally trigger the TLR 7/8 pathways. As the compound is introduced into the tumor, an immediate innate immune response is raised against the tumor tissues, likely through the release of inflammatory cytokines in the tumor microenvironment, which starts repolarizing immunosuppressive cells.^16–18, 34–36^ As the compound spreads throughout the tumor and the lymphatic system for an extended duration, a robust adaptive immune response is likely raised against the tumor. However, it is currently not possible to determine whether this response is driven solely by the unspecific sustained exposure to the TLR7/8a or if AASTY and its nanodisc-forming properties play a role by extracting and presenting tumor antigens in nanodiscs, or a combination of both. In either case, this approach appears to limit side-effects of systemic circulation of TLR 7/8 agonists, while yielding a robust therapeutic response, despite being a monotherapy. A combination therapy with complementary immune activators, can be expected to further strengthen this treatment.

## Conclusion

Poly(acrylic acid-co-styrene) was found to have prolonged tissue retention, when injected subcutaneously. This was exploited to yield a localized immunotherapy using a copolymer drug conjugate consisting of a TLR 7/8 agonist covalently bound to poly(acrylic acid-co-styrene). This resulted in significant potentiation of the therapy when tested in a CT26 murine carcinoma model, compared to the unbound TLR 7/8 agonist. Multiple mechanisms may be at play, as the copolymer is known to lyse cells, and form nanodiscs with the cell membranes, which could in term lead to greater immunostimulation, and an adaptive response. All surviving mice had adaptive immunity when re-innoculated with CT26 cells. Lastly, the polymer was shown to distribute to lymphoid tissues, further corroborating that it plays a direct role in the adaptive immune response to the tumor.

## Acknowledgement

The author thanks The Independent Research Fund Denmark (0171-00081B) for the support of this work. C.L.M is supported by Schmidt Science Fellows, in partnership with the Rhodes Trust.

**Notes** The authors declare the following competing financial interest(s): A.A.A.A. and N.J.B are listed as inventors on a pending patent describing some of the technology covered in this manuscript. Subsets of the figures were created with BioRender.com.

