## Supplementary material for "Cancer Immunotherapy through Tissue Adhering Polymers"

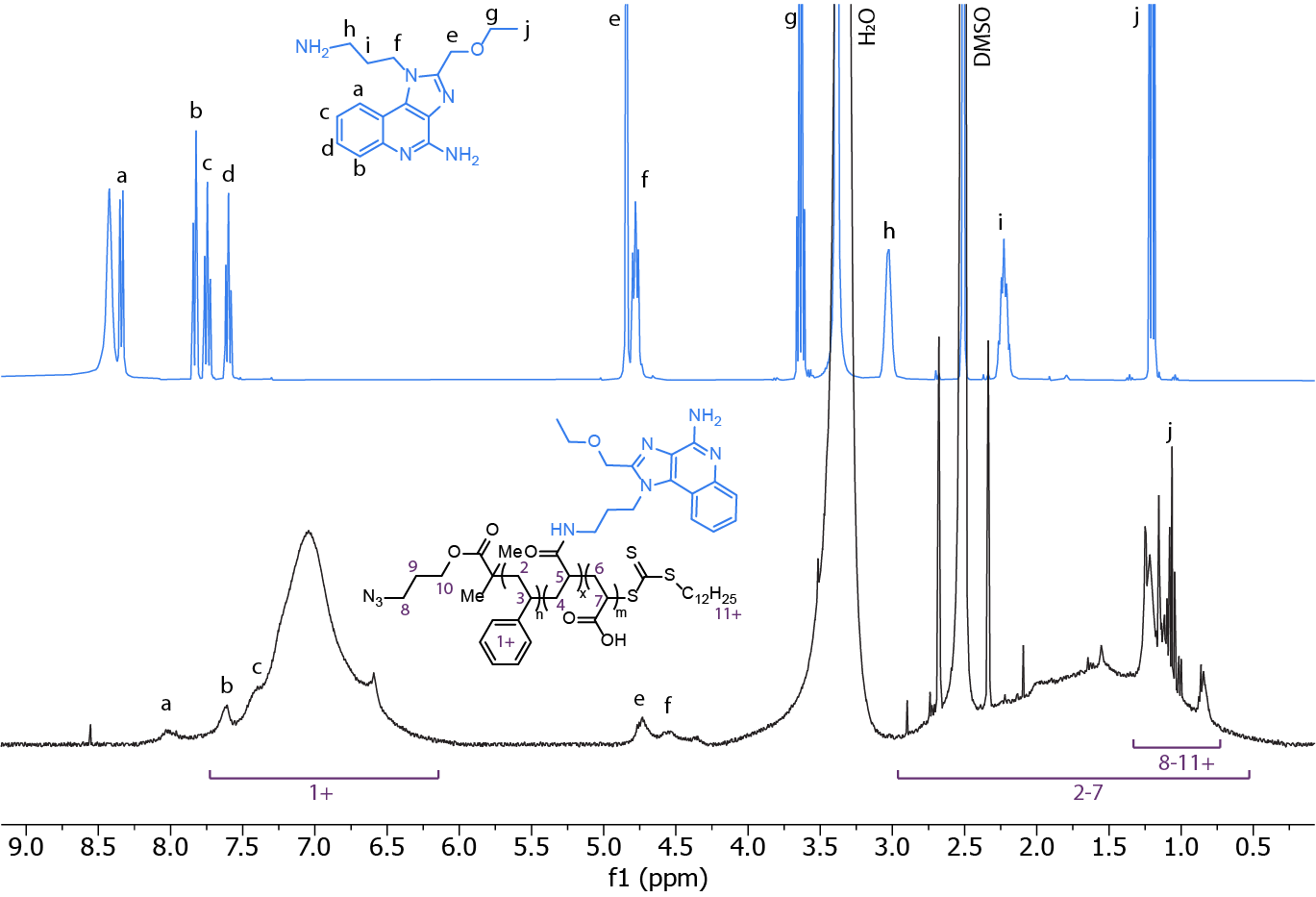

Figure S1: ^1^H NMR spectra of the free TLR7/8a and AASTY_9.9_-TLR7/8a.

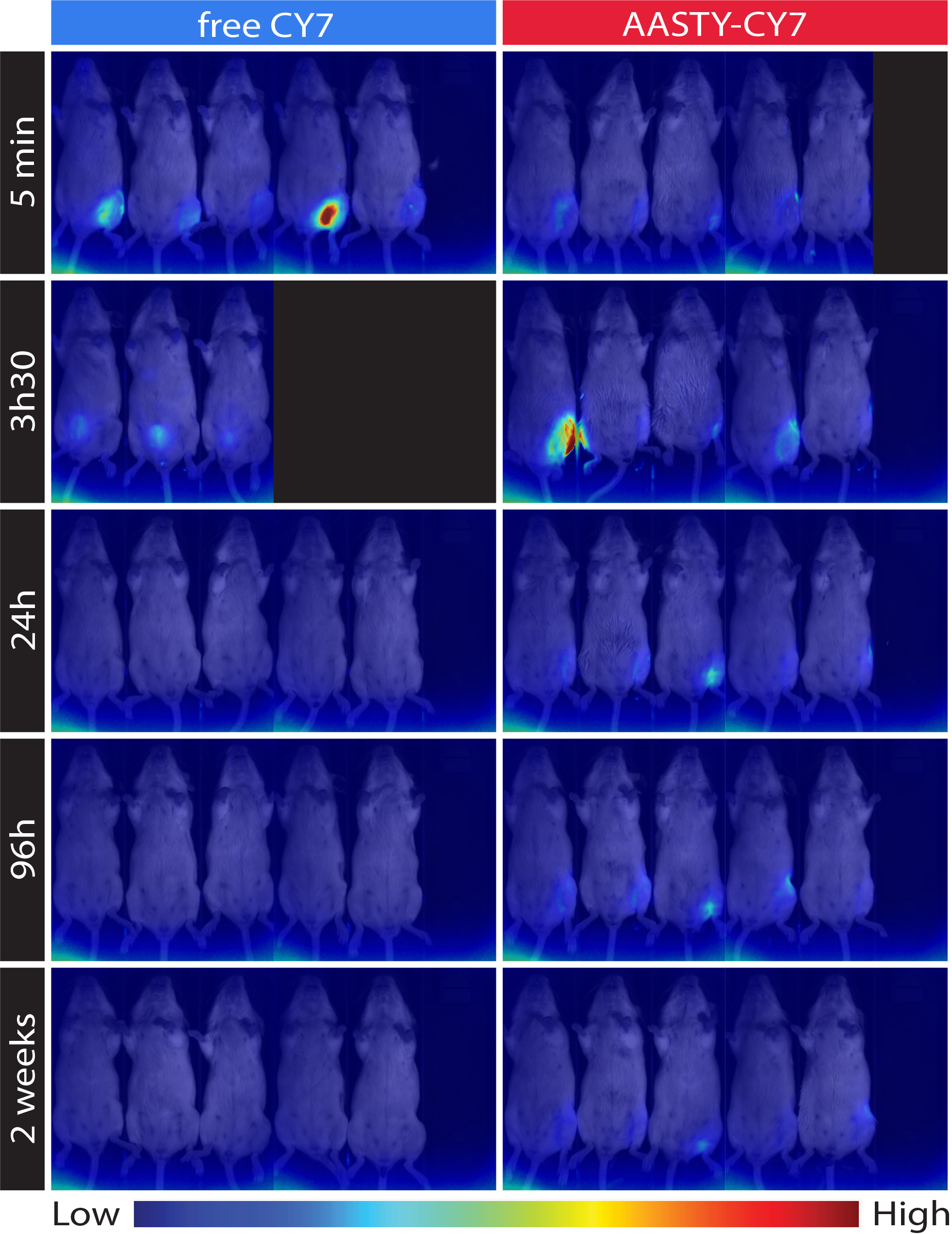

Figure S2: Additional biodistribution images of AASTY-Cy7 conjugates after SC injection. A rapid renal excretion and clearance into the bladder is observed for the water-soluble free Cy7 dye. The polymer conjugate on the other side remains at the injection point.

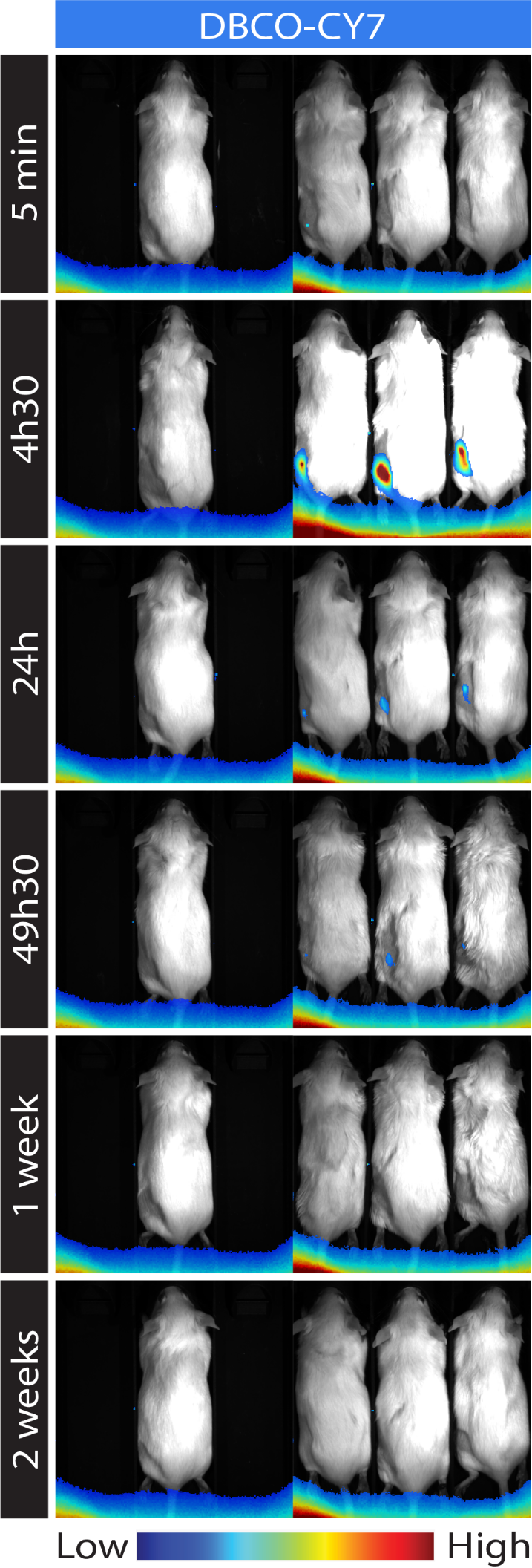

Figure S3: Control biodistribution experiment of DBCO-Cy7 in 5% DMSO-DPBS after SC injection. This Cy7 signal is hardly visible at the 5 min timepoint due to fluorescence quenching of the DBCO-dye at high concentration (this is also observed *in vitro*). After dilution in the SC tissue, the signal is then clearly visible after 4h30. Almost all the Cy7 signal then disappears within 1 week.

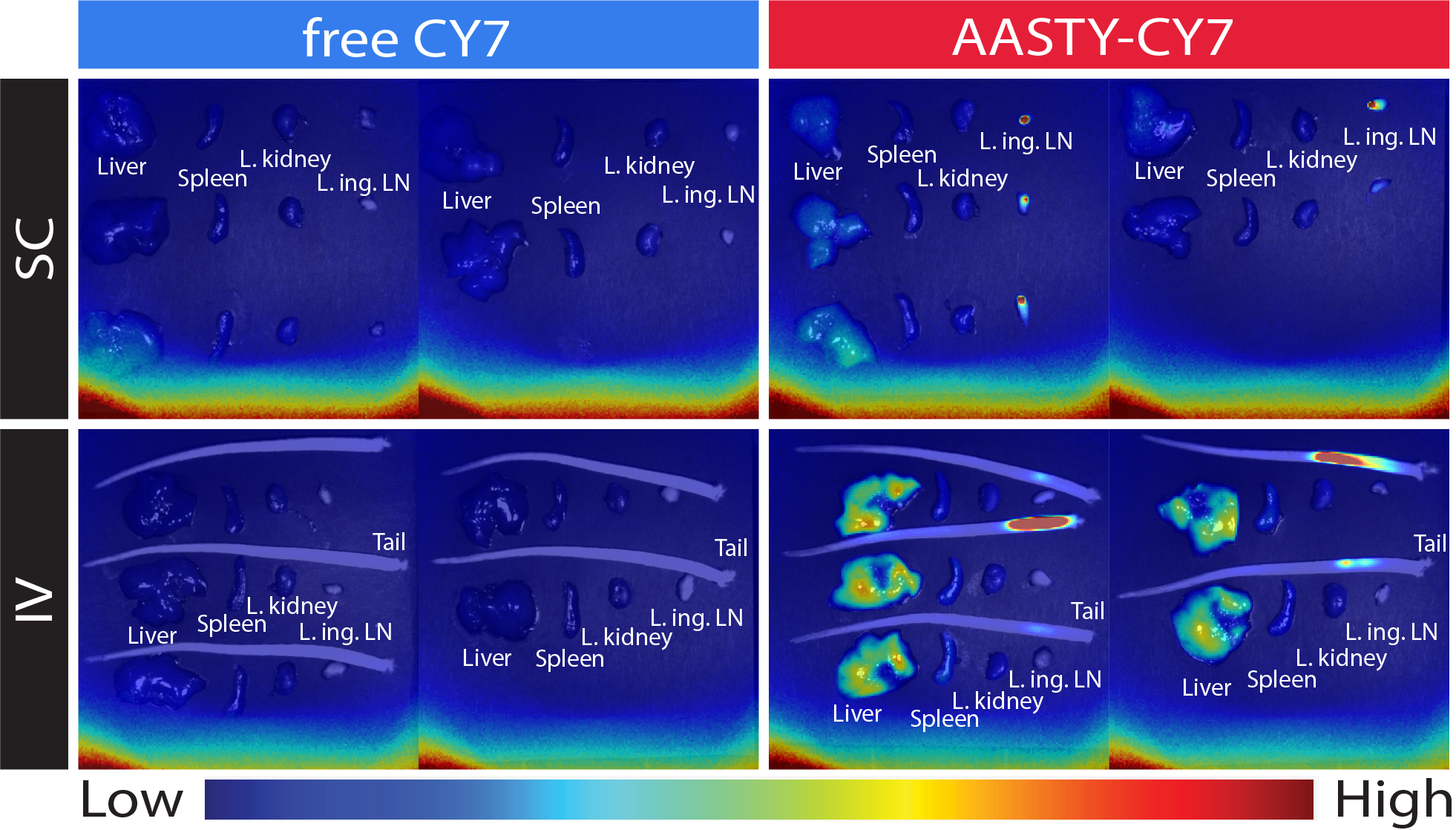

Figure S4: Biodistribution images of AASTY-Cy7 conjugates in isolated organs after SC and IV injection. The organs were immediately excised during autopsy, laid on a black cardboard sheet, and imaging was performed with the same imaging technique as *in vivo*. For IV administration, the tail was included in the imaging to account for the strong residual signal in the injection point. “L.” = left (injected side), “ing. LN” = inguinal lymph node.

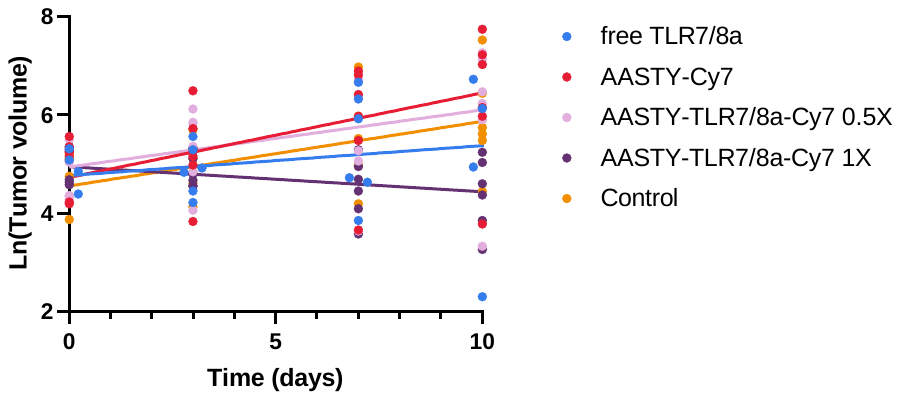

Figure S5: Log-transformed average tumor growth for the treatment groups for the first 10 days following administration of the first treatment dose. Tumor volumes were log-transformed to preserve normality of residuals and respect homoscedasticity. The curve parameters were estimated using a restricted maximum likelihood (REML) mixed model with mouse counting as random effect subject.

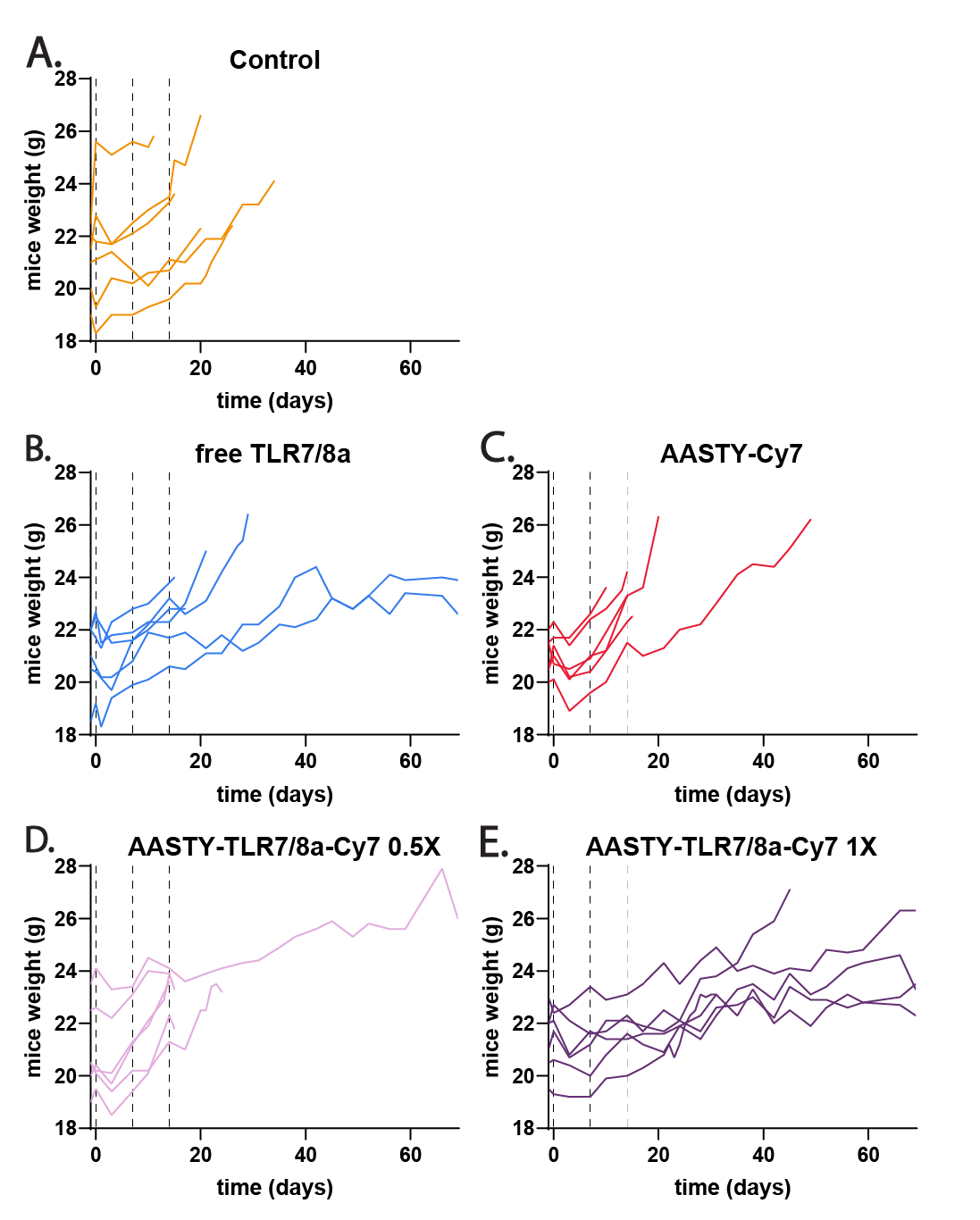

Figure S6: Individual mice weight curves over time. The dotted vertical lines represent the treatment injection time-points. None of the mice reached the end-point threshold of 10% weight loss overnight or 15% initial weight loss.

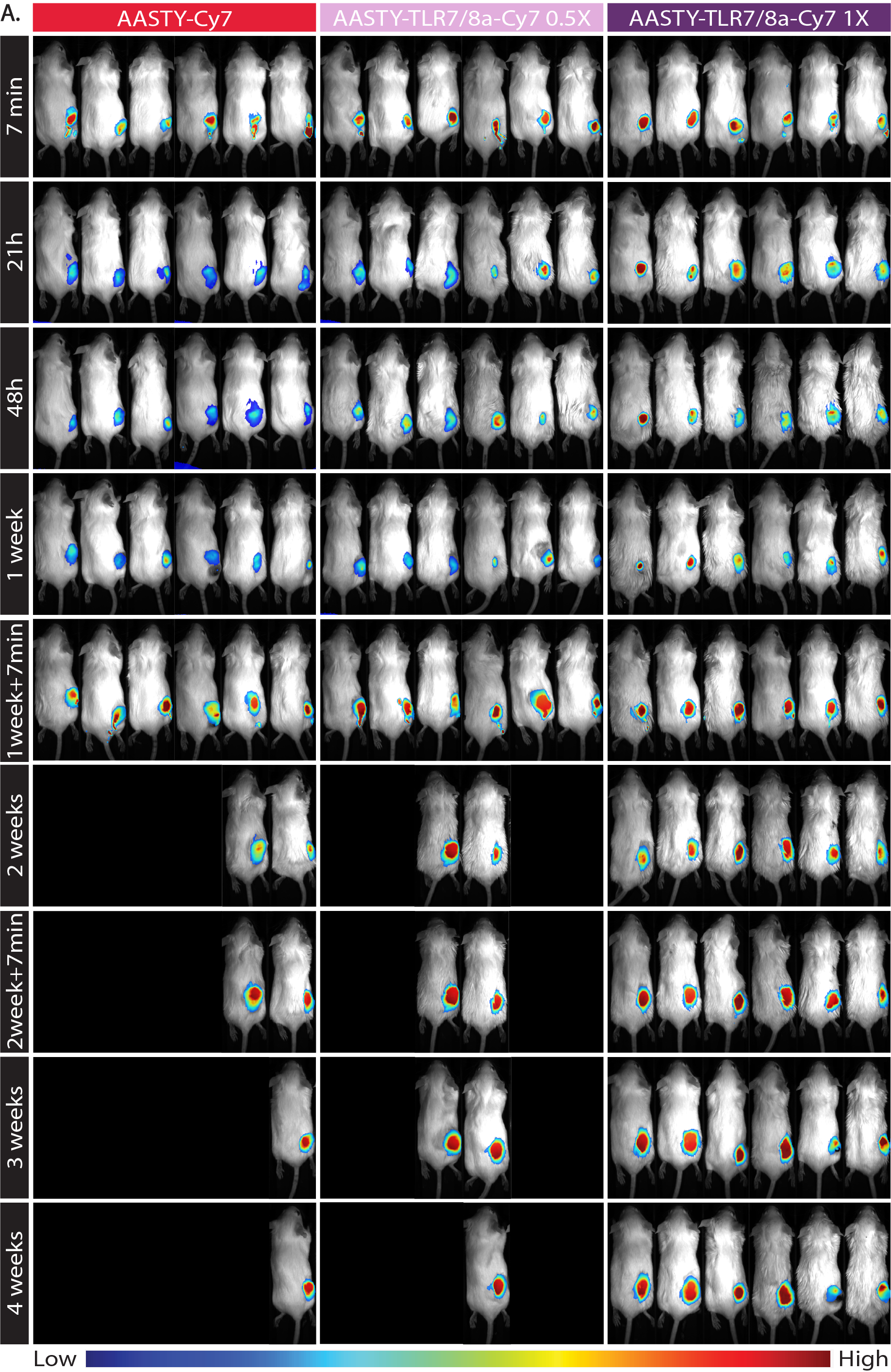

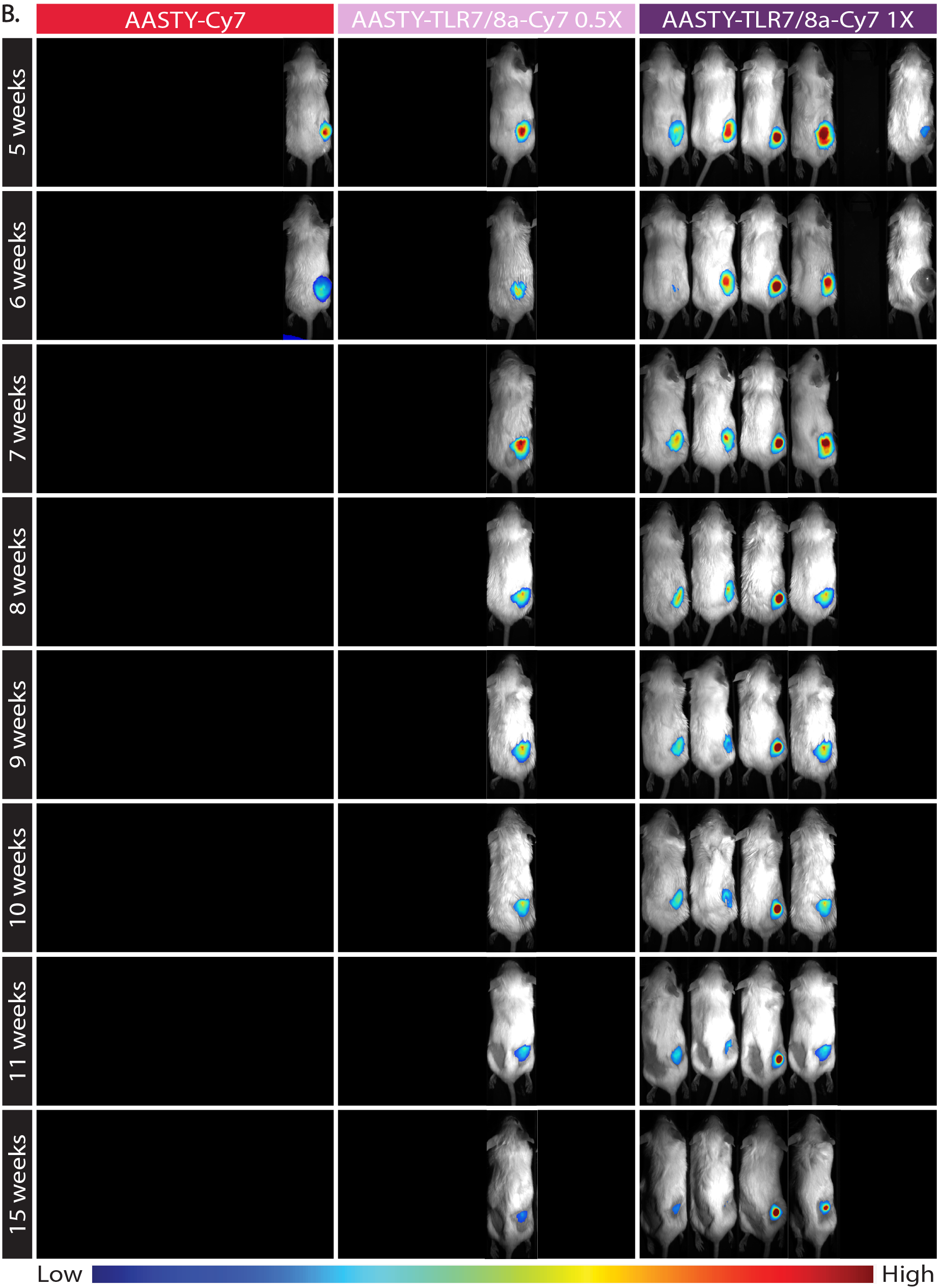

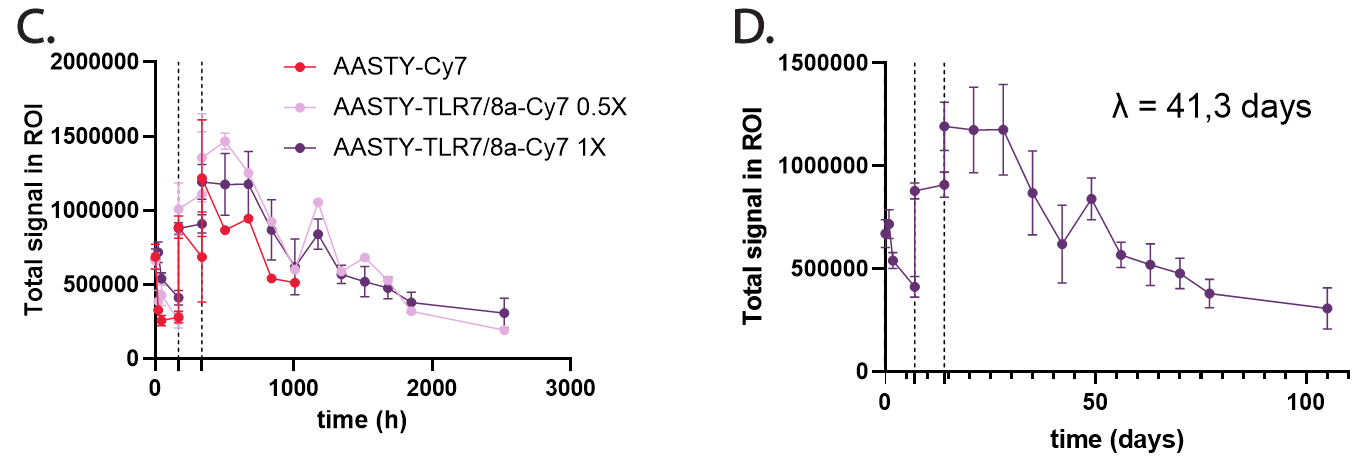

Figure S7: Additional biodistribution images of AASTY-TLR7/8a-Cy7 conjugates in tumors after IT injection (A: first 4 weeks; B: next 11 weeks) and Cy7 signal quantification (C: all samples; D only the AASTY-TLR7/8a-Cy7 1X group). (C,D) The dotted vertical lines represent the treatment injection time-points. (D) The half-life λ of the Cy7 conjugate in the tumor was estimated using a linear regression of the log-transformed integrated signal, as in the first biodistribution studies, on the data points following the third injection. For AASTY-Cy7, AASTY-TLR7/8a-Cy7 0.5X and AASTY-TLR7/8a-Cy7 1X respectively, n varies from 6 to 1, 6 to 1 and 6 to 4, based on the number of surviving mice at each timepoint.

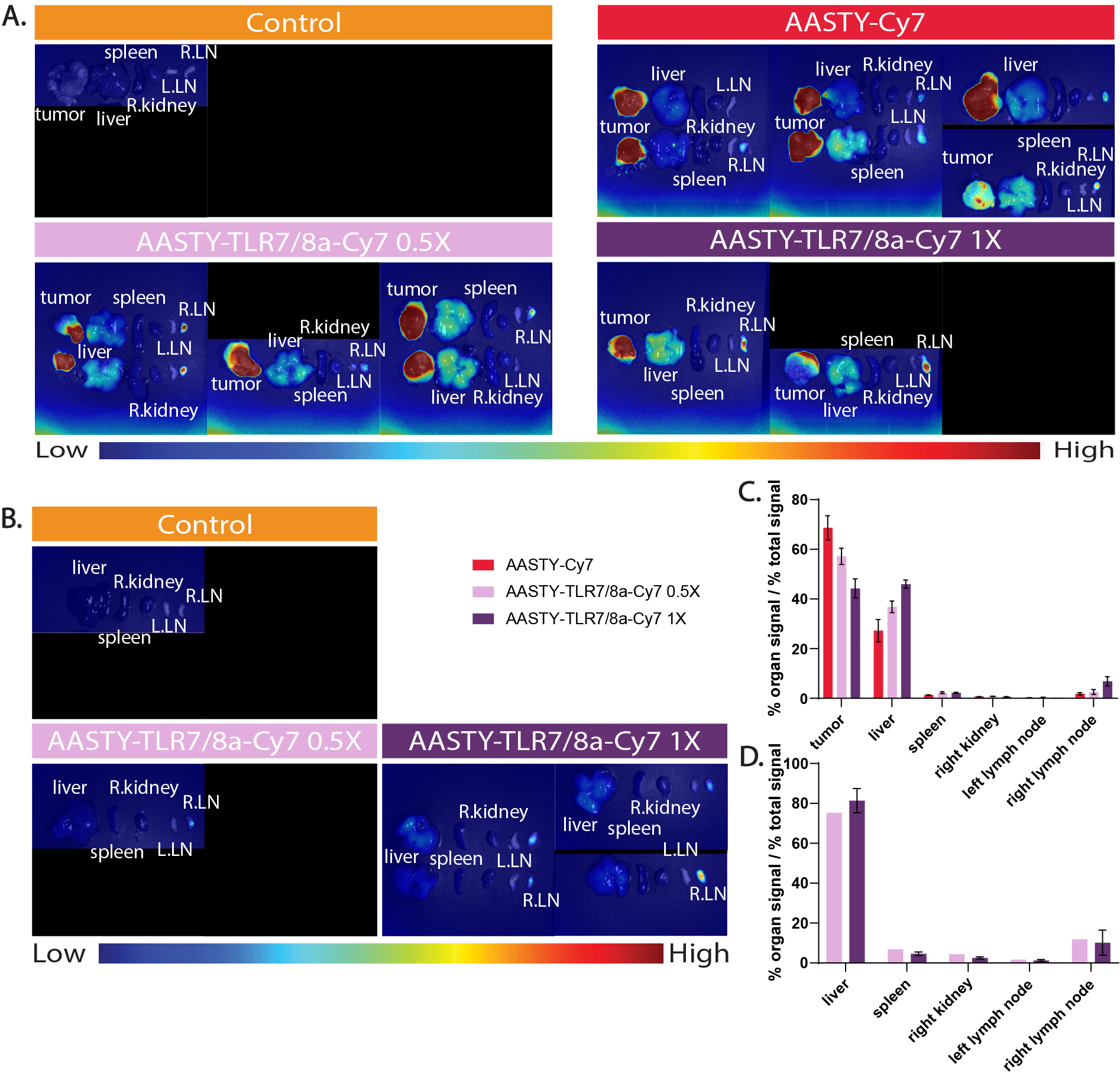

Figure S8: Biodistribution images of AASTY-TLR7/8a-Cy7 conjugates in isolated organs after IT injection (A,B) and Cy7 signal quantification (C,D). (A,C) represent the autopsied mice that were euthanized because they reached the tumor volume end-point (“uncured mice”). (B,D) represent the autopsied mice that were still alive at the end of the study (“cured mice”). The main difference between these 2 groups is the presence or absence of tumor at the point of autopsy. (A,B) “L. = left”, “R. = right” (injection side), “LN” = inguinal lymph node”. (C) For AASTY-Cy7, AASTY-TLR7/8a-Cy7 0.5X and AASTY-TLR7/8a-Cy7 1X respectively, n = 6, 5 and 2. (D) For AASTY-TLR7/8a-Cy7 0.5X and AASTY-TLR7/8a-Cy7 1X respectively, n = 1 and 4.
